# Endothelial-pericyte interactions regulate angiogenesis via VEGFR2 signaling during retinal development and disease

**DOI:** 10.1101/2025.03.08.642174

**Authors:** Ying-Yu Lin, Emily Warren, Bria L. Macklin, Lucas Ramirez, Sharon Gerecht

## Abstract

Pericytes stabilize the microvasculature by enhancing endothelial barrier integrity, resulting in functional networks. During retinal development, pericyte recruitment is crucial for stabilizing nascent angiogenic vasculature. However, in adulthood, disrupted endothelial-pericyte interactions lead to vascular dropout and pathological angiogenesis in ocular microvascular diseases, and strategies to stabilize the retinal vasculature are lacking. We demonstrate that direct endothelial-pericyte contact downregulates pVEGFR2 in endothelial cells, which enhances pericyte migration and promotes endothelial cell barrier function. Intravitreal injection of a VEGFR2 inhibitor in mouse models of the developing retina and oxygen-induced retinopathy increased pericyte recruitment and aided vascular stability. The VEGFR2 inhibitor further rescued ischemic retinopathy by enhancing vascularization and tissue growth while reducing vascular permeability. Our findings offer a druggable target to support the growth of functional and mature microvasculature in ocular microvascular diseases and tissue regeneration overall.

## Introduction

The vascular system is responsible for delivering nutrients and oxygen to the body as well as facilitating communication between organs. It develops from the differentiation of endothelial cells (ECs) from angioblasts(Majesky, 2018), and is the first functioning physiological system that forms during embryogenesis. Following the formation of the initial vasculature, ECs then proliferate, migrate, and sprout to extend vascular networks to support organ function. In humans, the retinal vasculature extends from the optic nerve head, growing radially and penetrating through the retinal layers to form functional vascular networks(Gariano & Gardner, 2005). The recruitment of mural cells, such as pericytes to the capillaries and microvasculature, and smooth muscle cells to the larger vessels, results in matured and functional blood vessels that are capable of transporting oxygen and nutrients. Vascularization of injured, diseased, or engineered tissue is required for tissue healing and regeneration, and eventual homeostasis (Masson-Meyers & Tayebi, 2021; Veith *et al*, 2019; Yang *et al*, 2020).

The vascular endothelial growth factor (VEGF)/VEGF Receptor 2 (VEGFR2) signaling pathway is essential in forming organ vasculature(Karaman *et al*, 2022). VEGFR2 is activated by VEGF-A, -C, and -D(Apte *et al*, 2019) and plays an important role during angiogenesis(Wang *et al*, 2020). The establishment of ligand-receptor dimerization upon VEGF binding leads to the phosphorylation of the downstream tyrosine residues of VEGFR2 at the catalytic tyrosine kinase domain (TKD) and the activation of signaling pathways that regulate EC function and vascular formation (Clegg & Mac Gabhann, 2015). Several phosphorylation sites are responsible for VEGFR2 activation and cell signaling regulation. The phosphorylation of tyrosine 951 promotes Akt activity to affect EC survival and permeability(Smith *et al*, 2020; Zhou *et al*, 2022). Additionally, phosphorylation of VEGFR2 at the surface of the tip cell mediates tip cell migration and increases glycolysis (Gerhardt *et al*, 2003) (De Bock *et al*, 2013; Yeh *et al*, 2008). It has been demonstrated that cancer cells secrete Angiopoietin-1 following VEGFR2 inhibition, leading to temporary pericyte recruitment and tumor vascular normalization(Winkler *et al*, 2004). However, it remains unknown if and how VEGFR2 activity in ECs regulates pericyte recruitment outside of the cancer microenvironment. Here, we sought to examine whether VEGFR2 in ECs modulates pericyte recruitment and endothelial barrier function during retina development and retinopathy.

Pericyte contact with the capillaries/microvasculature is critical for ECs to maintain barrier function and vascular homeostasis (Payne *et al*, 2020; Wang *et al*, 2014; Zhao & Chappell, 2019). Platelet-derived growth factor-BB (PDGF-BB) secreted by ECs has been shown to regulate pericyte recruitment, proliferation, and migration during vascular development (Payne *et al*., 2020). Pericytes guide vascular remodeling by migrating along sprouting ECs (Payne *et al*, 2019; Stapor *et al*, 2014) and secreting extracellular matrix (ECM) degradation molecules (Carmeliet & Jain, 2011). Pericytes also secrete ECM, which contributes to the vascular basement membrane to support organ function (Sakhneny *et al*, 2021; Yamazaki *et al*, 2020). Therefore, for successful tissue regeneration or healing, the re-vascularization process must include pericytes. However, despite the current understanding of the initial pericyte recruitment, it remains unclear how pericytes respond and modulate the initial shift from angiogenesis to stabilization in ECs.

Diabetic retinopathy is a microvascular disease characterized by pericyte loss, increased vascular permeability, and endothelial cell death, followed by the growth of abnormal blood vessels and vision impairment (Ansari *et al*, 2022). The available treatment options for revascularizing a damaged retina is limited to intravitreal injections of anti-angiogenic agents, corticosteroids and non-steroid anti-inflammatory drugs (Mansour *et al*, 2020; Wang & Lo, 2018). VEGF is a key driver of diabetic retinopathy disease progression(Caldwell *et al*, 2003), and is negatively correlated with pericyte recruitment(Schrimpf *et al*, 2014). Neutralizing excessive VEGF secretion is used to treat retinopathy, including FDA-approved drugs such as aflibercept (VEGF-trap), faricimab, and ranibizumab. However, these drugs primarily sequester VEGF-A, preventing their binding to all VEGFRs and impacting a wide range of VEGF downstream signaling, potentially making them less effective while leading to unwanted side effects and therapeutic resistance(Fallah *et al*, 2019; Sharma *et al*, 2023). Furthermore, not all patients respond effectively to treatments, and methods to reverse vascular damage are yet to be developed (Simó & Hernández, 2022; Whitehead *et al*, 2018). Pericyte destruction is the first pathological event observed in diabetic retinopathy mice (Hammes *et al*, 2004), followed by the switching of ECs to an angiogenic phenotype (Watson *et al*, 2017). Therefore, it is important to consider pericytes’ role in modulating angiogenesis through cellular crosstalk. However, the mechanisms of revascularizing damaged tissue or re-establishing EC-pericyte communication in diseases such as diabetic retinopathy, have remained largely unidentified (Park *et al*, 2017; van Splunder *et al*, 2023).

Here we set out to determine the role of pericytes in regulating VEGFR2 signaling on ECs, and subsequent vascular stabilization and maturation in both developing and diseased retina. Using isogenic ECs and pericytes derived from human induced pluripotent stem cells (hiPSCs), we begin by examining the hypothesis that after initial pericyte recruitment by PDGF-BB (Kemp *et al*, 2020), a direct physical contact between pericytes and ECs is required to downregulate VEGFR2 activity in ECs. Next, using hiPSC *in vitro* modeling and *in vivo* mouse models of the oxygen-induced retinopathy and the healthy developing retina, we examine the hypothesis that the downregulation of VEGFR2 activity amplifies pericyte recruitment to the nascent vascular networks, leading to network stabilization. Using a specific VEGFR2 inhibitor, we demonstrate that downregulating VEGFR2 pY951 restores endothelial barrier function and vascular stability by enhancing pericyte recruitment to the developing and diseased retina. This result informs the potential of blocking VEGFR2 activity to treat injured and diseased tissue or to vascularize tissue-engineered organs by stabilizing developing vasculature through increased pericyte recruitment.

## Results

### hiPSC-derived pericytes express characteristic markers and responsiveness to PDGF-BB

To better understand EC-pericyte interactions, we employ hiPSCs to co-differentiate into a bicellular population of ECs and pericytes (Cho *et al*, 2020; Kusuma *et al*, 2013; Smith *et al*, 2018). We used glycogen synthase kinase (GSK) 3 inhibitor CHIR99021 to induce mesoderm followed by subsequent differentiation into CD31^+^ ECs (iECs) based on an established protocol (Chan *et al*, 2021; Macklin *et al*, 2022). hiPSC-derived pericytes (iPericytes) were obtained by culturing the CD31^-^ cells in the pericyte growth media for five days (**Fig 1A**). Simultaneously obtaining iECs and iPericytes from the same hiPSC line allows for a shared genetic background that is ideal for cell-cell interaction studies.

**Fig. 1.**
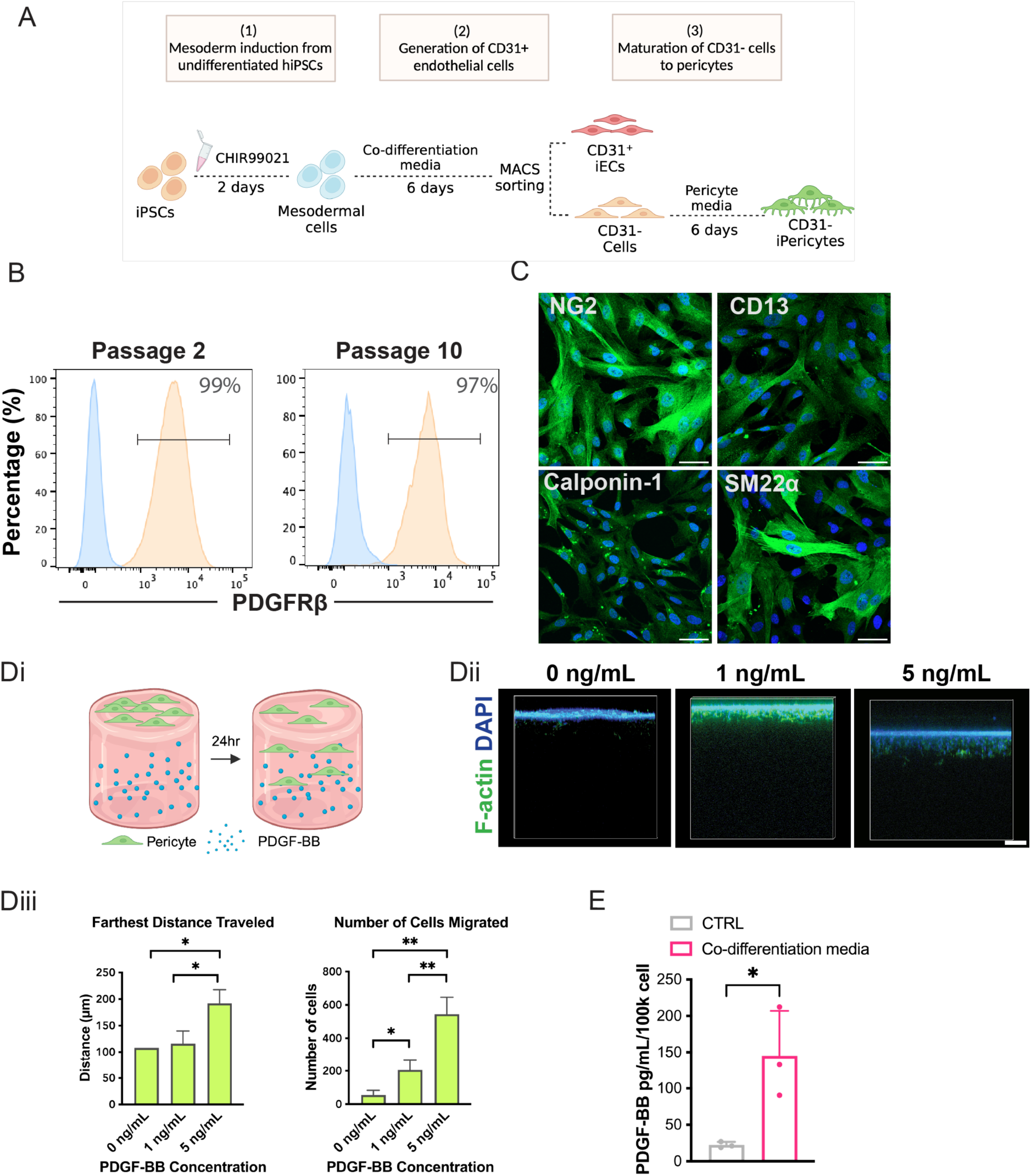
Characterization of iPericytes. (**A**) Schematic illustration of iPericyte differentiation. (**B**) Flow cytometry analysis of PDGFRb expression in iPericytes. (**C**) Immunofluorescence images of pericyte markers at passage 2 post sort: NG2, calponin-1, CD13, and SM22α (all in green; nuclei in blue). Scale bar: 50µm. (**D**) i. Schematic illustration of iPericyte recruitment experiment; ii-iii. Representative confocal images showing side views of hydrogels with pericytes (F-actin in green, nuclei in blue) and corresponding quantification is showing dose-dependent iPericyte migration in response to PDGF-BB. Bars represent means±SD; N=2 (0 ng/ml), N=3 (1ng/ml, 5ng/ml). Scale bar: 50µm. (**E**) ELISA analysis of PDGF-BB levels in the media. Illustrations created with BioRender.com. Bars represent means±SD; Significance levels were set at *\*p* ≤ 0.05, and *\*\*p* ≤ 0.01.

The derived CD31^+^ population (iECs) was expanded and characterized with immunofluorescence staining for EC markers, including PECAM-1(CD31), vascular endothelial cadherin (VE-cadherin), and von Willebrand factor (vWF) (**Fig S1**). The identity of iPericytes was confirmed by flow cytometry analysis and immunofluorescence staining. The iPericytes expressed pericyte markers, including platelet-derived growth factor receptor beta (PDGFRβ), neural glial antigen-2 (NG2), CD13, calponin-1, and SM22α (**Fig 1B-C**). Expression was maintained for up to 10 passages (**Fig S2A**). It is worth noting that PDGFRβ is consistently expressed for extended culture, implying their ability to maintain phenotype through passages (**Fig S2B**).

Next, we evaluated if iPericytes have functional PDGFRβ receptors, as PDGF-BB/PDGFRβ signaling is essential for initial pericyte recruitment to ECs (Smyth *et al*, 2022). We simulated pericyte recruitment by seeding iPericytes on the top of type I collagen hydrogels with or without exogenous PDGF-BB in the gel for 24 hours (**Fig 1Di**). Notably, iPericytes migrate toward the bottom of the hydrogel containing PDGF-BB in a dose-dependent manner (**Fig 1Dii-iii**), demonstrating the functionality of the highly expressed of PDGFRβ in iPericytes.

To confirm that iECs can recruit iPericytes, we examined PDGF-BB secretion. Here, PDGF-BB secretion from iECs cultured in ECGM supplemented with VEGF and TGF-β inhibitor (control) was compared to iECs cultured in co-differentiation conditioned media collected prior to sorting (ECGM supplemented with VEGF and TGF-β inhibitor). Surprisingly, iECs secrete more PDGF-BB when cultured in the co-differentiation conditioned media (**Fig 1E**). Therefore, we determined that the presence of iPericytes leads to an increase in PDGF-BB secretion from iECs.

### VEGFR2 inhibition in iECs leads to increased pericyte recruitment

In a previous study, we found that iECs are highly angiogenic with abundant expression of VEGFR2 (Macklin *et al*., 2022). When vessels are in an angiogenic state, they are highly migratory with increased junctional permeability allowing for extensive remodeling (Schwartz *et al*, 2018). Furthermore, VEGFR2 is negatively correlated with pericyte recruitment (Greenberg *et al*, 2008). We thus hypothesized that pericyte recruitment and vessel stabilization require downregulation of VEGFR2, which is achieved through direct physical contact between ECs and pericytes.

To investigate whether modulating VEGFR2 signaling could affect the ability of ECs to recruit pericytes, we began by treating iECs with VEGFR2 inhibitor ZM323881. First, we analyzed EC-derived factors that have been reported to be important for pericyte migration, invasion, and proliferation (Kemp *et al*., 2020). After 48 hours of VEGFR2 inhibitor treatment, qRT-PCR was performed on iECs. We found upregulation of PDGF-BB, heparin-binding EGF-like growth factor, and Endothelin-1 when compared to the group without inhibitor treatment (**Fig 2A**).

**Fig. 2.**
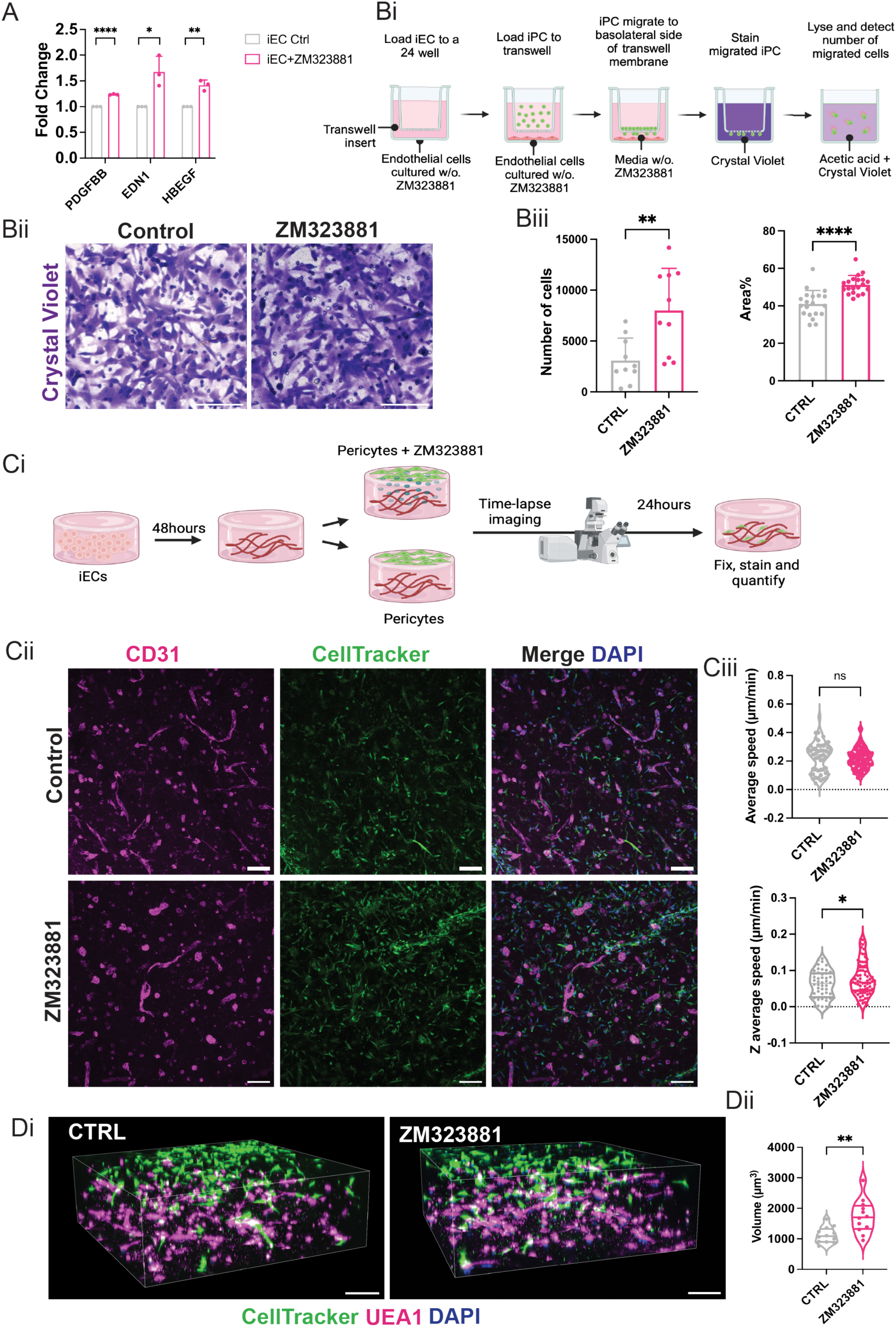
Inhibition of VEGFR2 in iECs enhances iPericyte recruitment. (**A**) RT-qPCR for PDGF-BB, heparin-binding EGF-like growth factor (HB-EGF), and Endothelin-1 (EDN1) in iECs treated with VEGFR2 inhibitor ZM323881 for 48 hours. Bars represent means±SD; N=3. (**B**) i. Schematic illustration of 2D migration assay; ii. Representative images of migrated iPericytes stained with crystal violet. Scale bar: 300 µm. iii. Quantification of the percentage of area covered by cells and the number of cells migrated. N=3, total of 10 samples for each group with two fields of view for each sample. (**C**) i. Schematic illustration of iPericytes recruitment to 3D vascular networks; ii. Representative immunofluorescence images of iECs networks (CD31, magenta) and iPericytes (CellTracker, green; Nuclei, blue) at 24 hours. Scale bar: 100 µm. iii. Quantification of average iPericyte migration speed and average migration speed in the Z axis; n= 6 - 7 gels for each group, total of 134 cells were analyzed. (**D**) i. Representative immunofluorescence images of iECs networks (UEA1, magenta) and iPericytes under hypoxia (CellTracker, green; Nuclei, blue) at 24 hours. Scale bar: 150 µm. ii. Quantification of iPericyte volume in the gel; 4 gels from each group with three fields of view for each sample. Illustrations created with BioRender.com. Bars represent means±SD; Significance levels were set at not significant (ns) *p* > 0.05, *\*p* ≤ 0.05, *\*\*p* ≤ 0.01, and *\*\*\*\*p* ≤ 0.0001.

Next, we asked if we would observe increased pericyte recruitment by inhibiting VEGFR2. Using a Transwell migration assay, we found that VEGFR2 inhibition increased iPericyte migration with increased cell coverage and cell number (**Fig 2B**).

We then asked if we could enhance pericyte recruitment to 3D vascular networks by inhibiting VEGFR2 signaling. To model pericyte migration to endothelial networks, iECs were encapsulated in the type I collagen hydrogels for 48 hours where they self-assembled into 3D vascular networks. iPericytes were then seeded on the top of the hydrogel-containing vascular networks with or without a VEGFR2 inhibitor to mimic pericyte recruitment during capillary network formation (**Fig 2Ci**). We observed increased pericyte recruitment in the inhibitor-treated group where more CellTracker positive cells were recruited to the vascular networks (**Fig 2Cii**). We also examined iPericyte migration speed, and while we found no differences in the average cell migration speed across experimental groups, we observed increased cell migration speed of iPericytes in the Z axis in the treatment group compared to the control group (**Fig 2Ciii**). These data suggest that VEGFR2 inhibition leads to increased iPericyte migration and recruitment through iEC chemoattraction mechanisms.

Finally, to recapitulate the pathological environment in diabetic retinopathy, iPericytes were seeded on top of the hydrogel-containing iEC vascular networks under hypoxic conditions. Here, too, we observed greater pericyte penetration, evident by an increase in the total volume of pericytes in the VEGFR2 inhibitor-treated group compared to the control (**Fig 2D**).

Overall, the in vitro results indicate that the inhibition of VEGFR2 enhances pericyte recruitment to the vascular networks, potentially through the upregulation and increased secretion of pericyte recruitment factors, leading to increased EC-pericyte interactions.

### Pericytes modulate pVEGFR2 activity via direct EC contact

We next sought to determine whether pericytes regulate VEGFR2 expression through soluble factors or through direct contact. We first examined VEGFR2 pY951 because it is one of the major phosphorylation sites of VEGFR2, and has been shown to be the key regulator of EC survival, migration, and permeability during angiogenesis (Wang *et al*., 2020). We cultured iECs in iPericyte-conditioned media (**Fig 3A**) or co-cultured iECs with iPericytes (**Fig 3D**). Western blotting was conducted to analyze VEGFR2 activity and compared among treatment groups over the course of four days. We found that VEGFR2 pY951 levels in iECs did not change along the culture with iPericyte-conditioned media (**Fig 3B-C**). In contrast, the expression of VEGFR2 pY951 in iECs was decreased along the co-culture with iPericytes for three to four days (**Fig 3E-F**). The same expression pattern was observed with another VEGFR2 phosphorylation site Y1175, which also participated in regulating vascular permeability (**Fig S3**). This suggests that pericytes modulate EC VEGFR2 activity only by establishing direct physical contact with ECs. We confirmed the absence of VEGFR2 in both iPericytes and primary human retinal pericytes, regardless of VEGF supplementation in the culture media (**Fig S4**). These results confirm that measured VEGFR2 activity is exclusive to the ECs.

**Fig. 3.**
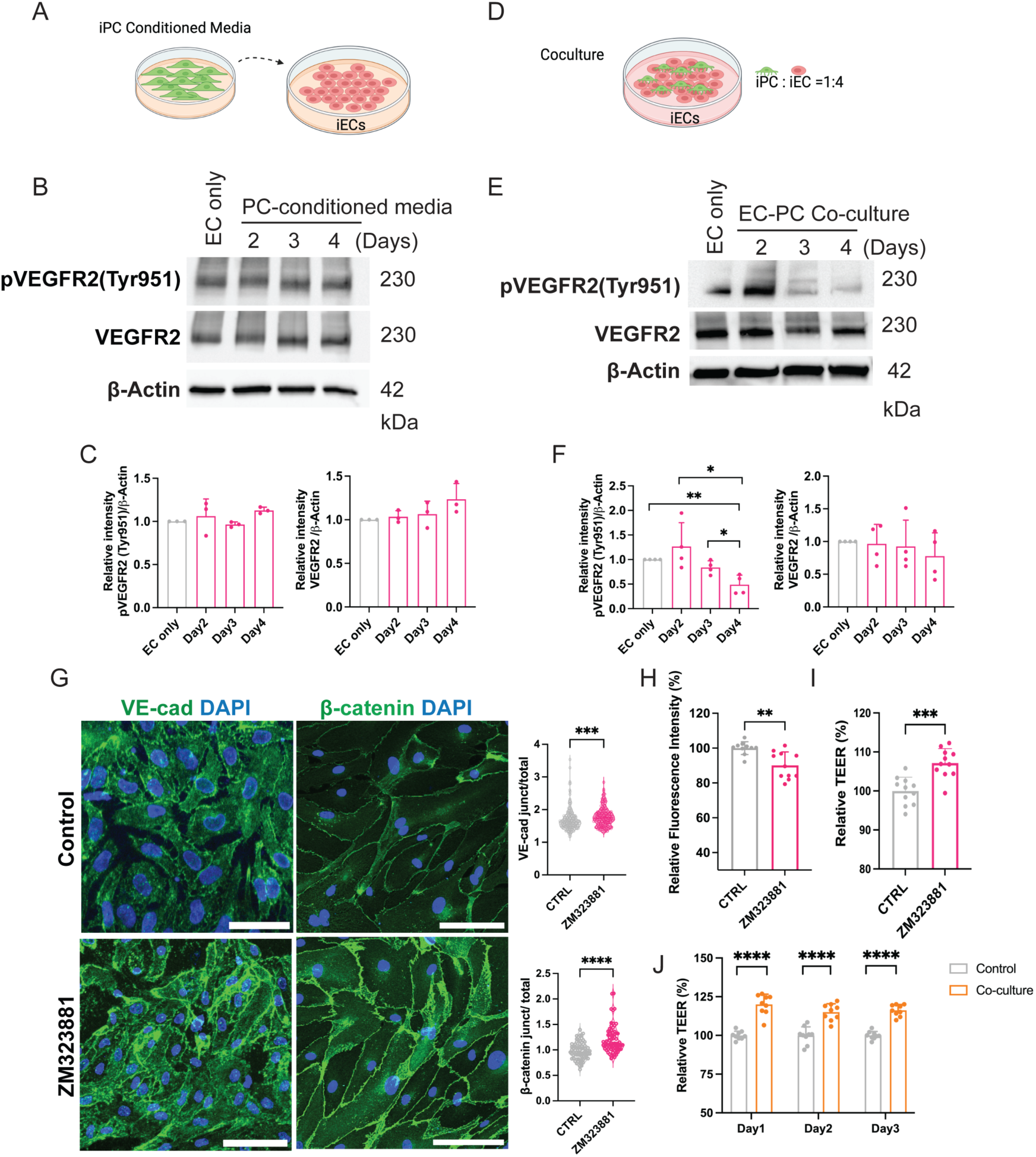
Direct EC-pericyte contact is required for VEGFR2 pY951 downregulation, leading to barrier stabilization. (**A**,**D**) Schematic illustrations of conditioned media and co-culture experiments. (**B**,**E**) Representative western blots for VEGFR2 pY951, VEGFR2, and beta-actin of control iEC and after cultured in iPericyte-conditioned media or iEC-pericyte co-culture for two, three, and four days. (**C**,**F**) Quantification of western blot for pVEGFR2 pY951 and VEGFR2 normalized to beta-actin loading control and vehicle control. Bars represent means±SD N=3. (**G**) Representative max intensity projection of immunofluorescence images of VE-cadherin, and β-catenin in iECs treated with or without VEGFR2 inhibitor for 24 hours. Quantification of junctional intensity of VE-cadherin, and β-catenin. N=3, with two to five fields of view for each sample. Scale bar: 50 µm. (**H**) FITC-dextran Transwell permeability assay on iECs treated with or without VEGFR2 inhibitor for 24 hours. Bars represent means±SD; N=3. (I) TEER measurements on iECs treated with or without VEGFR2 inhibitor for 24 hours. N=3 (J) TEER measurements on iECs co-cultured with or without iPericytes. N=3. Illustrations created with BioRender.com. Bars represent means±SD; Significance levels were set at *\*p* ≤ 0.05, *\*\*p* ≤ 0.01, *\*\*\*p* ≤ 0.001 and *\*\*\*\*p* ≤ 0.0001.

VE-cadherin forms adherens junctions between ECs that are important in regulating vascular permeability (Claesson-Welsh *et al*, 2021). When ECs are in the angiogenic state, they exhibit increased vascular permeability allowing sprouting and lumen formation (Felmeden *et al*, 2003; Krüger-Genge *et al*, 2019). Once new vessels are formed, ECs become quiescent, and the vascular permeability decreases. Decreased vascular permeability indicates that the vasculature is stabilized, becoming functional and mature. To examine whether inhibition of VEGFR2 signaling enhances VE-cadherin adherens junctions and barrier function, iECs were treated with VEGFR2 inhibitor. As expected, inhibiting VEGFR2 resulted in the downregulation of VEGFR2 pY951 at both early and later time points (**Fig S5**). Interestingly, inhibition of VEGFR2 pY951 did not significantly impact VE-cadherin expression level but rather increased VE-cadherin and β-catenin localization to the cell membrane of iECs (**Fig S5; Fig 3G**). To examine the impact on barrier function, we analyzed cell permeability using fluorescein isothiocyanate-Dextran (FITC-Dextran) and trans-endothelial electrical resistance (TEER) using EVOM3. We found decreased cell permeability (**Fig 3H**) and increased TEER (**Fig 3I**) when VEGFR2 is inhibited in iECs. The co-culture of pericyte and iECs on the Transwell membranes results in an increase in TEER compared to the control group (**Fig 3J**). Overall, these results indicate that inhibition of VEGFR2, such as through pericyte direct contact, increases vascular barrier function contributing to vascular network stabilization.

### VEGFR2 inhibition increases pericyte recruitment in the developing mouse retina

Next, we sought to test if the role of VEGFR2 is critical in EC-pericyte interactions during developmental angiogenesis. To achieve this, healthy pups were used as a developing mouse model to examine how VEGFR2 inhibitors impact pericyte recruitment. It has been established that the retinal vessels and pericytes start to emerge, grow, and penetrate the intermediate and deep layer of the retina from around postnatal day 7 onwards (Darche *et al*, 2022; Stahl *et al*, 2010). Thus, we performed an Intravitreal injection on postnatal day 8 (**Fig 4A**). Staining with pericyte marker NG2, revealed that pericyte coverage is increased in the intermediate and deep plexus in the healthy retina treated with VEGFR2 inhibitor compared with untreated retinas (**Fig 4B**). Our findings demonstrate that inhibiting VEGFR2 could strengthen pericyte recruitment to the endothelial vasculature, aiding in vascular stability during developmental angiogenesis. Overall, our data presents a previously unknown mechanism that governs pericyte-mediated vascular stabilization, in which downregulation of VEGFR2 occurs through direct contact between ECs and pericytes. This interaction results in enhanced pericyte recruitment, ultimately leading to the stabilization of the nascent vasculature (**Fig 4C**).

**Fig. 4.**
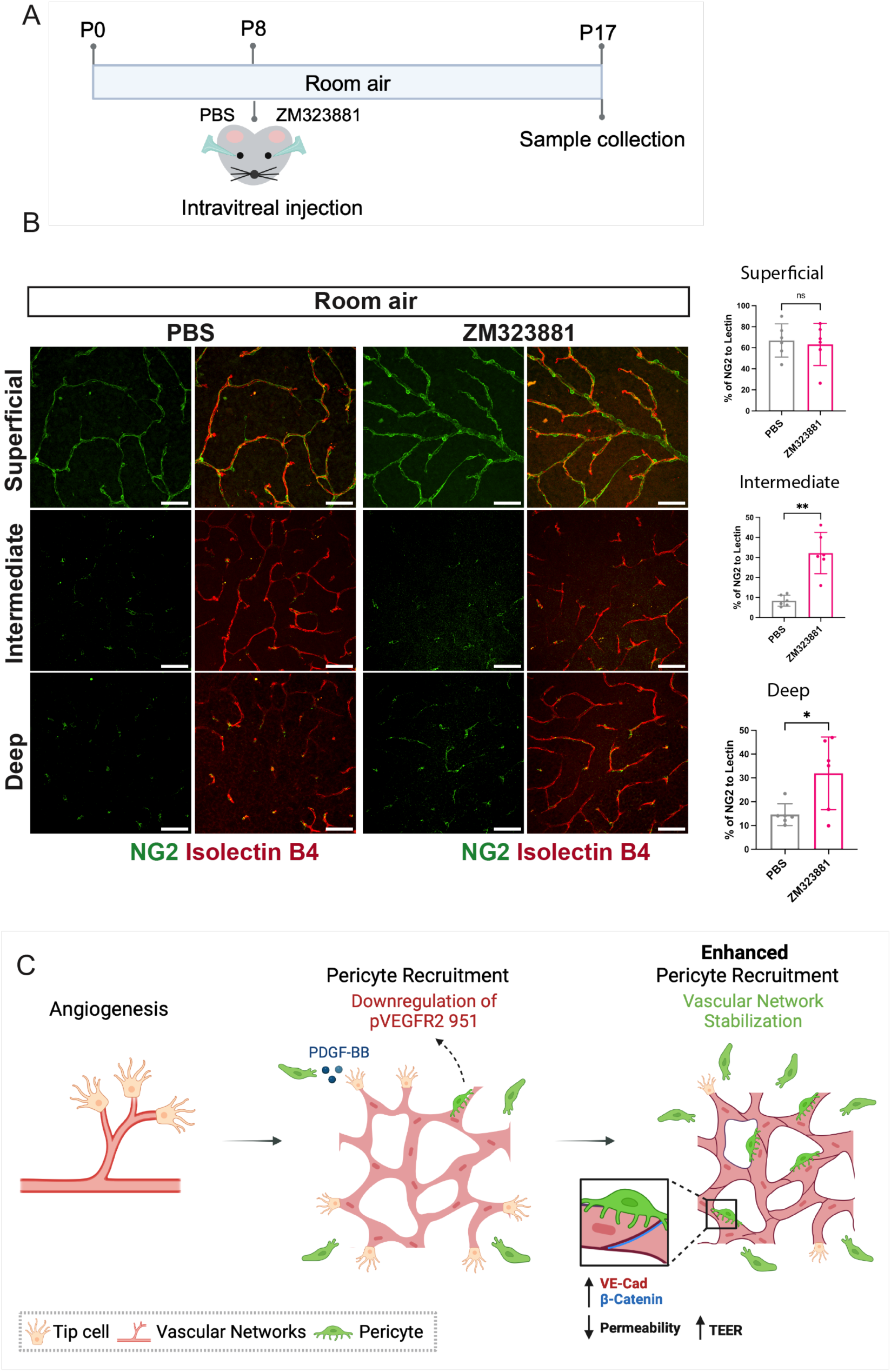
Pericyte recruitment is increased in the developing mouse retina treated with VEGFR2 inhibitor. (**A**) Schematic illustration of experimental schedule in the developing mouse. (**B**) ***Left*** - Representative immunofluorescence images of the healthy mouse retinal tissue at P17 throughout different layers. ***Right*** - Quantification of the presence of pericytes (NG2, green) relative to mouse vasculature (Isolectin B4, red) (n=6). Scale bar: 50µm. (**C**) Conclusion schematic showing that downregulation of VEGFR2 through direct contact between ECs and pericytes enhances pericyte recruitment, thereby leading to the stabilization of the nascent vasculature. Illustrations created with BioRender.com.

### Inhibition of VEGFR2 enhances pericyte recruitment, resulting in improved healing in the OIR mouse model

To examine the translation implications of our findings, we sought to investigate if inhibiting VEGFR2 *in vivo* could restore vascular stability in ischemic retinopathy by enhancing pericyte recruitment. Therefore, we chose the oxygen induced retinopathy (OIR) mouse model to study pericyte recruitment under pathological angiogenesis. Intravitreal injection of VEGFR2 inhibitor or vehicle control PBS was performed on postnatal day 12 when pups were returned to room air after 5 days of exposure to 75% oxygen to induce ischemic retinopathy (**Fig 5A**). On P17, we confirmed the downregulation of VEGFR2 pY951 in the inhibitor-injected retinas (**Fig 5B**). We found an increase in revascularization through a decrease in pathological neovascularization (NV) and less vaso-obliteration (VO) in the VEGFR2 inhibitor-treated OIR retina when compared to the contralateral eye injected with vehicle control PBS (**Fig 5C**). To confirm the specificity of the VEGFR2 inhibition, we examined the efficiency of an FDA-approved pan-VEGFR inhibitor, Tivozanib, to treat the OIR mouse. Tivozanib was approved by FDA in 2021 to treat relapsed or refractory advanced renal cell carcinoma(Beckermann *et al*, 2024). We found no significant decrease in VO and a significant reduction of NV in the inhibitor-treated retina (**Fig S6A**). Given the critical role of VEGF in neovascularization in the OIR model (Vähätupa *et al*, 2020), these data suggest that the downregulation of VEGFR2 activity ameliorates pathological angiogenesis under hypoxia.

**Fig. 5.**
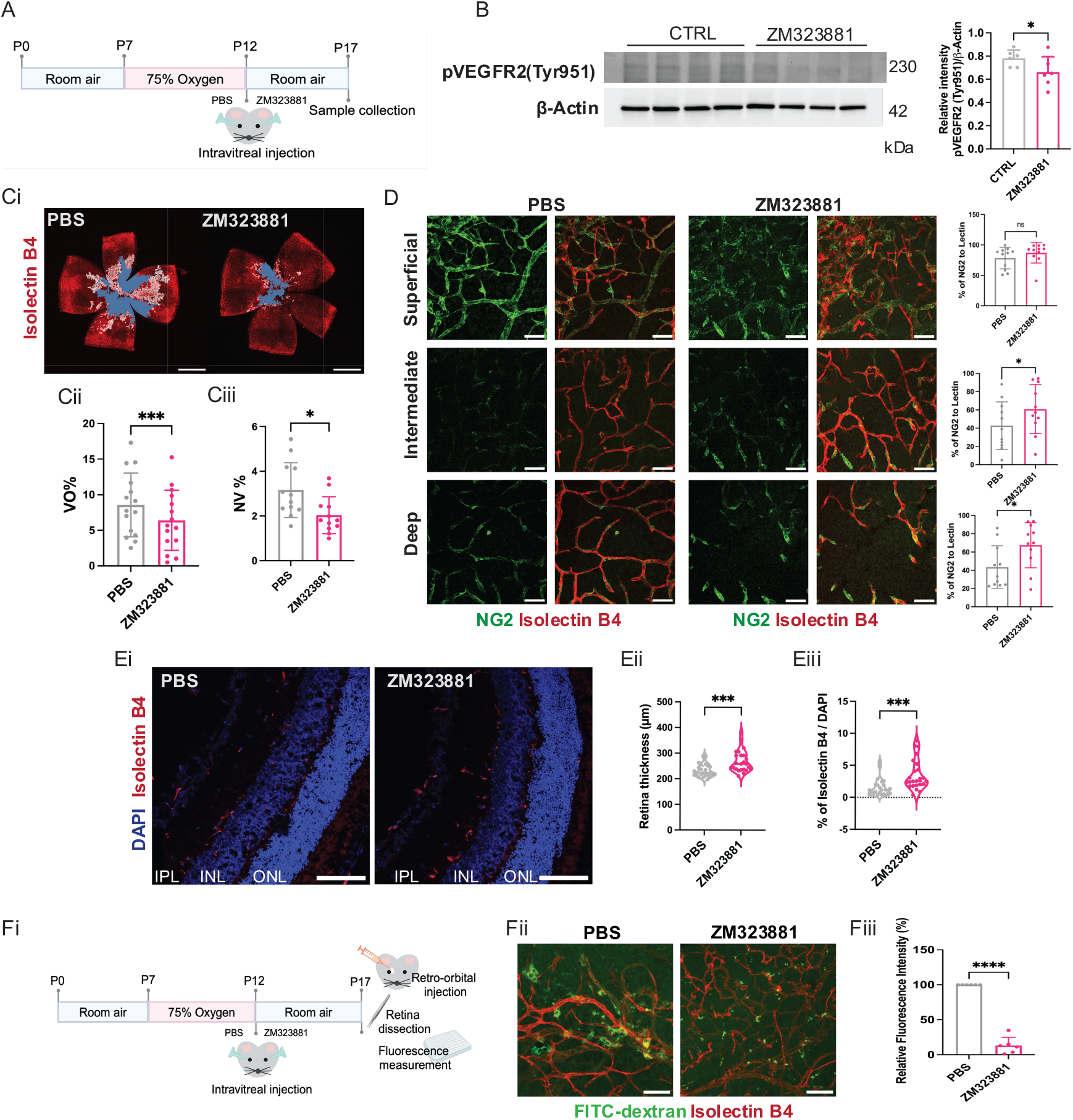
Downregulation of VEGFR2 improves retinal healing following OIR by enhanced pericyte recruitment and vascular function. (**A**) Schematic illustration of experimental schedule of OIR mouse model. (**B**) Representative western blots for VEGFR2 pY951 and beta-actin in mouse retina collated 24 hours after ZM323881 injection. (**C**) **i.** Representative immunofluorescence images of the OIR mouse retinal tissue at P17, five days after intravitreal injection of VEGFR2 inhibitor (Isolectin B4, red; VO area, blue; NV area, white). **ii-iii.** Quantifications of vaso-obliteration area (VO) and pathological neovascularization (NV) at P17 (n=15). Scale bar: 300 µm. (**D**) *Left* - Representative immunofluorescence images of the OIR mouse retinal tissue at P17 throughout different layers. *Right* - Quantification of the presence of pericytes (NG2, green) relative to mouse vasculature (Isolectin B4, red) (n=11). Scale bar: 50 µm. (**E**) **i.** Representative immunofluorescence images of the OIR mouse retina cross-section at P17. (**ii-iii**) Quantification of retina thickness and the percentage of vessel coverage (Isolectin B4, red) relative to retina area (DAPI, blue) (n=3 mice, 8 sections each). Scale bar: 100 µm. (**F**) **i**. Schematic illustration of experimental schedule of vascular permeability assay with OIR mouse model. **ii.** Representative immunofluorescence images of the OIR mouse retinal tissue (left). Scale bar: 50 µm. **iii.** Relative fluorescence intensity of FITC-dextran in OIR retinas treated with or without VEGFR2 inhibitor. Bars represent means±SD; n=6. Illustrations created with BioRender.com. Bars represent means±SD; Significance levels were set at not significant (ns) *p* > 0.05, *\*p* ≤ 0.05, *\*\*\*p* ≤ 0.001 and *\*\*\*\*p* ≤ 0.0001.

We next assessed the involvement of pericytes in ischemic angiogenesis in the OIR retina treated with or without VEGFR2 inhibitor by staining the retina with the pericyte marker NG2. The spatial development of mouse retinal vasculature is formed starting from the superficial to the intermediate and the deep plexus (Stahl *et al*., 2010). We thus analyzed pericyte recruitment to the vascular plexus in the three layers. As the superficial plexus vasculature is fully formed at postnatal day 17 (Stahl *et al*., 2010), we found no difference in pericyte coverage between the treated and control groups (**Fig 5D, left**). However, we found that the percentage of pericyte presence in the intermediate and deep layers of the retina is significantly increased in the VEGFR2 inhibitor-treated eyes compared to untreated controls (**Fig 5D**). In contrast, treatment with the pan-VEGFR inhibitor Tivozanib, increased pericyte recruitment in the superficial and intermediate but not in the deep layer of the retina compared to control (**Fig S6B**). The difference between the two treatments led us to suspect that pericyte-mediated vascular stabilization is mostly mediated by VEGFR2 rather than other VEGFRs.

Next, we stained the cross-sections of the retina from the OIR mice to further investigate if increased pericyte recruitment results in a healthier retinal structure. We observed a significant increase in retinal thickness in the VEGFR2 inhibitor group (**Fig 5Ei-ii**), with increased vessel coverage area when VEGFR2 is downregulated (**Fig 5Eiii**), suggesting that augmented pericyte involvement improves retina structure and vascular normalization.

Finally, to determine the impact of VEGFR2 inhibition on vessel functionality, we performed a vascular permeability assay on retinas injected with PBS control or VEGFR2 inhibitor (**Fig 5Fi**). Following FITC-dextran injection to the retro-orbital venous sinus, we found that retinal vascular leakage was reduced in the VEGFR2 inhibitor-treated retina as indicated by fluorescence imaging and direct quantification of the extracted FITC-dextran (**Fig 5Fii-iii**). This result again supports the role of VEGFR2 in vascular stabilization during pathological angiogenesis. Overall, these outcomes demonstrate that inhibition of VEGFR2 leads to a healthier and thicker retina, as evidenced by enhanced pericyte recruitment and reduced vascular permeability in an OIR mouse model.

Taken together, our findings demonstrate that inhibiting VEGFR2 could strengthen pericyte recruitment to the endothelial vasculature, aiding in vascular stability during pathological and physiological angiogenesis.

## Discussion

Pericytes play a critical role in stabilizing microvasculature and in maintaining microvascular homeostasis (Payne *et al*., 2020; van Splunder *et al*., 2023). In contrast, during vascular remodeling, pericyte detachment was observed in the embryonic dorsal skin (Payne *et al*, 2021) and loss of pericytes is reported in various microvascular diseases, such as diabetic retinopathy, Alzheimer’s disease, and stroke (Li & Fan, 2023). It has also been shown that low pericyte coverage on tumor vessels in cancer patients correlates with cancer metastasis (Teichert *et al*, 2017; van Splunder *et al*., 2023). Nonetheless, the mechanism by which pericyte stabilize vasculature in angiogenesis during the development of nascent vessels or during pathological angiogenesis is not fully understood.

Most *in vitro* studies utilize primary or immortalized cells from different origins. In this study, isogenic ECs and pericytes derived from the same hiPSC origin were used to analyze cell-cell interactions during vascular network formation. PDGF-BB is secreted by ECs to recruit pericytes to blood vessels (Payne *et al*., 2020). Consistent with this, we found that iPericytes migrate in response to PDGF-BB. Furthermore, iECs secrete more PDGF-BB when cultured in conditioned media collected from iEC and iPericyte co-differentiation compared to iECs expansion alone. VEGFR2 has been shown to be highly expressed in angiogenic ECs(Wang *et al*., 2020) and is essential in vasculogenesis and angiogenesis (Zarkada *et al*, 2015), (Macklin *et al*., 2022). Since VEGFR2 was reported to be negatively correlated with pericyte recruitment(Greenberg *et al*., 2008), we postulated that the increase in PDGF-BB secretion is controlled by the interactions between pericytes and VEGFR2. We inhibited VEGFR2 in iECs and observed the upregulation of pericyte recruitment-related genes PDGF-BB, HB-EGF, and EDN1 in iECs. It’s worth noting that HB-EGF, in addition to PDGF-BB, has been found to be secreted by ECs to attract mural cells during vascular network formation (Stratman *et al*, 2010) and EDN1 has been linked to pericyte survival and proliferation (Yamagishi *et al*, 1993).

We next examined the recruitment of pericytes to ECs and observed increased recruitment when VEGFR2 was inhibited. Our observation aligns with a seminal cancer study, which demonstrated that the blockage of VEGFR2 temporarily facilitates pericyte recruitment to the tumor vasculature(Winkler *et al*., 2004). The study concluded that pericyte recruitment is regulated by increased secretion of tumor cell-derived Ang-1. Building on this finding, our work uniquely shows that ECs themselves regulate pericyte recruitment through VEGFR2 modulation, which is particularly relevant in treating diabetic retinopathy. When treated with the inhibitor, there was a noticeable increase in iPericyte migration speed towards 3D vascular networks in the Z axis. However, there was no significant increase in the overall iPericyte migration speed in all three axes. This confirms that increased iPericytes migration during EC VEGFR2 inhibition was due to chemoattractants rather than randomly. In the Transwell migration assay, increased iPericyte migration was observed in the VEGFR2 inhibitor-treated group. This confirms that pericyte recruitment increases in developing EC tube networks when VEGFR2 activity is reduced (Bowers *et al*, 2020).

A previous *in vivo* study demonstrated that vessels lacking VEGFR2 pY951 expression in the murine embryonic body are mostly covered by pericyte-like cells while pericytes are missing from vessels containing VEGFR2 pY951(Matsumoto *et al*, 2005). Indeed, we found that pericytes modulate vascular stability through down-regulation of VEGFR2 pY951. Interestingly, we only observed down-regulation of pVEGFR2 in the co-culture group but not in the conditioned media-treated group. This result agrees with recent studies that demonstrate the formation of direct EC-pericyte cell-cell contact is crucial in vasculogenesis and endothelial cell maturation (Ayloo *et al*, 2022; Payne *et al*, 2022a). Therefore, we conclude that pericytes can regulate VEGFR2 signaling only by establishing a direct contact with ECs but not through paracrine signaling. Similar to VEGFR2 pY951 which modulates vascular permeability through the TSAd-Src-PI3K-Akt pathway(Matsumoto *et al*., 2005), other VEGFR2 phosphorylation sites, including Y1175 and Y801, have been implicated in regulating vascular permeability via the PLCγ-PKC pathway(Blanes *et al*, 2007; Sjöberg *et al*, 2023). Down-regulation of VEGFR2 pY1175 was also observed in the co-culture group. Although further studies are needed to elucidate the impact of pY1175 on EC function, this observation further confirmed the role of pericyte in EC stabilization.

VEGFR2 pY951 has also been shown to be responsible for the formation of EC adherens junctions in a knocked-out mouse model (Li *et al*, 2016). When we treated iECs with VEGFR2 inhibitor, adherens junction formation as well as barrier function were augmented, suggesting a more stable EC phenotype. We speculate that proteins mediating EC-pericyte cell signaling, such as N-cadherin or connexin-43, which require physical adjacency for their binding, mediate the dephosphorylation of VEGFR2 pY951 through downstream signaling pathways upon their interaction. N-cadherin adherens junctions form between ECs and pericytes and regulate the endothelial barrier by inducing VE-cadherin recruitment to EC junctions(Kruse *et al*, 2018). Furthermore, EC-pericyte gap junctions mediated by connexin-43 allow for the exchange of small molecules and ions that is critical during blood vessel formation(Payne *et al*, 2022b). Future experiments examining the involvement of these proteins in VEGFR2 downregulation upon EC-pericyte contact and their role in potential therapeutic resistance are warranted.

VEGFA-VEGFR2 and pericytes synergistically regulate blood vessel development in the mouse neonatal retina (Fruttiger, 2002). Depletion of pericytes in the early stage of vascular development results in elevated VEGFR2 expression, increased EC proliferation and sprouting (Eilken *et al*, 2017). We inhibited VEGFR2 in healthy pups and observed enhanced pericyte coverage in the retina. This finding is in agreement with a previous study that showed that pericyte coverage is reinstated in VEGFR2 inhibitor-treated mouse patent blood vessels (Greenberg *et al*., 2008). Our results provide a deeper understanding of the mechanism in which this occurs.

Pericyte loss is a common occurrence in ocular microvascular diseases, and no effective treatments have been developed to restore the interaction between endothelial cells and pericytes for the revascularization of damaged tissues. We observed increased pericyte coverage in the intermediate and deep plexus of the OIR retina when treated with VEGFR2 inhibitor. Retina thickness and vessel density are noticeably increased, while vascular permeability is decreased in the OIR eyes injected with VEGFR2 inhibitor. These findings indicate a sign of vascular normalization and, consequently, a healthier retina compared to untreated OIR eyes. Although it is arguable that reduced VO and NV in the mouse OIR retina observed in our study might be due to the result of specifically downregulating VEGFA-VEGFR2 interactions. However, other factors, including insulin-like growth factor 1 (Upreti *et al*, 2022), erythropoietin (Chen *et al*, 2008), and angiopoietin-2 (Hackett *et al*, 2002) have been suggested in aiding pathological angiogenesis in OIR mouse model. It is conceivable that pericytes may also contribute to the decrease of NV.

In addition to uncontrolled pathological angiogenesis, the loss of therapeutic efficacy following anti-VEGF treatment is a frequent challenge in managing DR. Acquired resistance and local adverse effects are two major resistant events that happen after anti-VEGF treatment. Unlike in cancer, therapeutic resistance in DR is primarily driven by hyperglycemia, chronic inflammation, and persistent vascular leakage(Rezzola *et al*, 2020; Semeraro *et al*, 2015). Given the limited role of alternative angiogenic pathways in DR(Khan *et al*, 2020), stabilizing leaky pathological vasculature is key to improving therapeutic outcomes(Al-Kharashi, 2018). Our study showed that VEGFR2 inhibition enhanced pericyte recruitment, which results in vascular normalization, therefore might decrease the chance of resistance often occurring following anti-VEGF treatment.

Above all, this research highlights the connection between pericytes and VEGFR2 regulation in ECs during vascular network stabilization in development and disease. We show that pericytes must establish direct cell-cell contact with ECs to downregulate VEGFR2 signaling. Then, the downregulation of VEGFR2 leads to increased pericyte recruitment, vessel stabilization, and improved vessel function. We demonstrated that targeting VEGFR2 signaling in retinopathy accelerates and augments retina healing, thus offering a therapeutic modality.

## Materials and Methods

### Maintenance and differentiation of hiPSCs

C1-2 hiPSCs were cultured on a Vitronectin-coated plate in Essential 8 medium (Thermo Fisher Scientific) with the media changed daily for 3 days. Endothelial differentiation was started with mesodermal induction when cells were 60-80% confluent by adding CHIR99021 (6 µM) (STEMCELL Technologies) to the Essential 6 medium (Thermo Fisher Scientific) with daily media changes. On day two of the mesodermal induction, cells were detached using TrypLE Express (Thermo Fisher Scientific) and seeded on a type I collagen coated plate at 2x10^4^ cells/cm^2^ with co-differentiation media containing Endothelial Cell Growth Media (ECGM; Promocell) supplemented with 10 µM SB-431542 (Cayman Chemical Company), 50 ng/mL VEGF-A (PeproTech), and 10 µM Y-27632 (Selleckchem). After 24 hours, the media was changed to ECGM supplemented with 10 µM SB-431542 (Cayman Chemical Company) and 50 ng/mL VEGF-A (PeproTech), following media change every other day for six days.

### Isolation of hiPSCs to endothelial cells and pericytes

On day eight of the differentiation, CD31^+^ cells were isolated with a magnetic-activated cell sorter (MACS; Miltenyi Biotech Bergisch Gladbach). Cells were washed with dPBS (Thermo Fisher Scientific) once and detached using TrypLE Express. After centrifuging, cells were then resuspended in 100 μL MACS buffer containing 0.5mM EDTA and 0.5% BSA in dPBS. 10 µL of PE-conjugated anti-human CD31 (BD Biosciences) were added to the MACS buffer, and cells were incubated at 4°C for 10 minutes. After incubation, cells were washed with MACS buffer twice to remove unbound primary antibodies. Next, cells were incubated with 20 µL of anti-PE microbeads (Miltenyi Biotec Bergisch Gladbach) and 80 µL of MACS buffer at 4°C for 15 minutes. Cells were washed once with MACS buffer and sorted using the MS MACS separation column (Miltenyi Biotec Bergisch Gladbach). The purified CD31^+^ cells were expanded in ECGM supplemented with 10 µM SB-431542 (Cayman Chemical Company), and 50 ng/mL VEGF-A (PeproTech) on a type I collagen-coated plate. CD31^-^ cells were expanded in Pericyte Medium (ScienCell) on a type I collagen-coated plate.

### VEGFR2 inhibition

For all 2D *in vitro* inhibition studies, cells were cultured in ECGM (Promocell) supplemented with 10 µM SB-431542 (Cayman Chemical Company) and 25 ng/ml VEGF-A (PeproTech). For all 3D *in vitro* inhibition studies, cells were cultured in ECGM (Promocell) supplemented with 50 ng/ml VEGF-A (PeproTech). ZM323881 (Selleckchem) was dissolved in DMSO and added to the cell culture media at 1 µM. Cells were incubated with the inhibitor for 0.5, 1, 12, 24, or 48 hours, as indicated. DMSO was used as vehicle control.

### Flow Cytometry analysis

Cells were harvested for analysis using TrypLE (Invitrogen) dissociation buffer and collected in 100 µL of 0.1% BSA. Cells were then incubated with primary antibody for 30 minutes on ice. Antibodies are detailed in **Table S1**. Cells were washed 3 times with 0.1% BSA and passed through a 40-µm cell strainer. Flow analysis was conducted on a BD FACSCanto flow cytometer. Following manufacturer instructions, dead cell populations were gated out with forward-side scatter plots. All analyses were conducted using FlowJo software.

### Immunofluorescence staining and imaging

iECs, iPericytes, and 3D constructs were fixed in 3.7% paraformaldehyde (Sigma-Aldrich) for 10 minutes or in ice-cold methanol for 5 minutes. Cells fixed with paraformaldehyde were permeabilized with 0.1% Triton X-100 (Sigma-Aldrich). Cells were washed and blocked in 1% BSA for one hour at room temperature or overnight at 4°C. Cells were incubated with primary antibodies (**Table S1**) overnight at 4°C. The next day, cells were washed three times with 0.1% TWEEN 20 (Sigma-Aldrich) in PBS. Cells were incubated with secondary antibodies for 1 hour at room temperature. Primary and secondary antibodies are detailed in **Table S1**. Cells were washed three times with 0.1% TWEEN 20 (Sigma-Aldrich) in PBS followed by DAPI staining for 10 minutes at room temperature. Samples were imaged on a Nikon AX-R Confocal microscope using Element software. Adherens junctions were quantified using FIJI/Image J (NIH). 8-10 cells per image from triplicates for each experiment condition were randomly selected for manual tracing along cell-cell junctions.

### Quantitative reverse transcriptase PCR gene analysis

Total RNA was extracted using TRIzol reagent (Thermo Fisher Scientific) and purified using the RNeasy Mini Kit (Qiagen). RNA quality and concentration were measured using a nanodrop spectrophotometer. Complementary DNA (cDNA) was generated using GoScript Reverse Transcriptase Random Primers kit (Promega) per the manufacturer’s protocol. The TaqMan Universal PCR Master Mix and Gene Expression Assay were used for the genes of interest. TaqMan PCR was performed using the QuantStudio 3 PCR System. The results were calculated as 2^-Δ^ ^Δ^ ^CT^ obtained by comparing the cycle threshold (CT) between samples as normalized to the endogenous control gene TATA-binding protein (TBP). **Table S1** lists all primers.

### Western blot protein analysis

Cells were lysed using RIPA Buffer (Thermo Fisher Scientific) with 1x Protease and Phosphatase Inhibitor Cocktail (Thermo Fisher Scientific). Retinas were collected 24 hours after intravitreal injection to assess VEGFR2 activity. Retinas were lysed using RIPA Buffer (Thermo Fisher Scientific) with 1x Protease and Phosphatase Inhibitor Cocktail (Thermo Fisher Scientific). The concentration of extracted protein was quantified using Pierce BCA Protein Assay Kits (Thermo Fisher Scientific). Twenty to thirty micrograms of protein lysate were loaded into a 4% to 20% Mini-PROTEAN TGX precast protein gel (Bio-Rad) with electrophoresis and then transferred to a PVDF membrane. Membranes were blocked with 5% milk or BSA for 1 hour and incubated in primary antibody (**Table S1**) overnight at 4°C. Primary antibodies were detected by using HRP-linked secondary antibody (Cell Signaling Technologies). Membranes were visualized with Clarity Western ECL Substrate (Bio-Rad) and imaged using Bio-rad ChemiDoc. Blots were analyzed using imaging J. Primary and secondary antibodies are detailed in **Table S1.**

### 3D PDGF-BB pericyte recruitment assay

Collagen gels were prepared as described below, with the addition of PDGF-BB (1 ng/mL, 5 ng/mL and DMSO as vehicle control). iPericytes (75,000 cells/well) were seeded on the top of polymerized collagen gels in ECGM (Promocell) and invasion was allowed to occur over 24 hours. After 24 hours, gels were fixed then stained for Phalloidin and DAPI. The migration of cells into gels was analyzed for the number of cells invading and the distance from the top using the IMARIS software Spots package.

### ELISA analysis of PDGF-BB

iECs were seeded at 13,000 cells/cm^2^ and cultured for 48 hours in ECGM (Promocell) supplement with 10 µM SB-431542 (Cayman Chemical Company), and 50 ng/mL VEGF-A (PeproTech) or conditioned media collected on Day 8 of endothelial cell differentiation. PDGF-BB secretion was quantified by enzyme-linked immunosorbent assay (ELISA; ThermoFisher Scientific) following the manufacture’s protocol. Data were normalized relative to 100,000 cells.

### Transwell migration assay

iECs were seeded on a type I collagen-coated 24-well plate and cultured in the incubator until confluency. iECs were then treated with VEGFR2 inhibitor or vehicle control for 24 hours. The next day, the basolateral side of the Transwell inserts with an 8 μM pore size (Corning 3464) were coated with type I collagen for 1 hour at room temperature. iPericytes were seeded on the apical side of the Transwell inserts with VEGFR2 inhibitor or vehicle control and cultured for 6 hours in the incubator. iPericytes that did not migrate through the filters were removed with cotton swabs. iPericytes migrated to the basolateral side of the Transwell insert were fixed with methanol and stained for 20 min with 0.2% crystal violet in distilled water. Images were taken using an EVOS microscope. The area of iPericytes coverage was quantified using ImageJ. For cell number quantification, iPericytes were lysed with 10% acetic acid and the optical density was analyzed using a microplate reader at 595nm. Cell density was quantified by fitting absorbance values into a standard curve. Standard curves were generated by seeding iPericytes at various cell densities into a 96-well plate for 24 hours. Cells were then fixed with methanol, stained with 0.2% crystal violet and lysed with 10% acetic acid. Optical density was analyzed using a microplate reader (BioTek Synergy, Agilent) by measuring the OD at 595nm. Three to four wells of Transwell inserts were used for each experiment with two fields of view for each sample.

### 3D collagen vascular network assay

Collagen gels were prepared as described previously (Kusuma *et al*, 2014). Briefly, Rat Tail Collagen-I (Corning) was diluted in 1X PBS to form an 8 mg/mL working solution. To prepare 1 mL of collagen gel solution, 0.8-2x10^6^ iECs were resuspended in 210.1 uL Medium 199(1X) (Gibco), 336.5 uL ECGM supplemented with 50 ng/mL VEGF, 38.4 uL Medium 199(10X) (Gibco), and 375 uL of collagen-I solution. The pH of the gel solution was adjusted by adding up to 40 uL 1 M NaOH. Then, 62.5 uL of the hydrogel solution was added to each well of a 96 well and polymerized at 37 °C for 30 minutes. ECGM supplemented with 50 ng/mL VEGF-A (PeproTech) was added to the gels after 30 minutes. Constructs were incubated for 48 hours to allow network formation with daily media change. For hypoxia experiments, hydrogel constructs were moved to a hypoxia chamber (1% O2) at 37°C after 24 hours and incubated for additional 24 hours. After a total of 48 hours of incubation, iPericytes labeled with CellTracker™ Green (Thermo), according to the manufacturer’s protocol, were passaged to the hydrogel networks (0.5-1.5x10^5^ cells/well) with ECGM supplemented with 50 ng/mL VEGF with or without VEGFR2 inhibitor. The hydrogel constructs were returned to the incubator or hypoxia chamber and media with or without VEGFR2 inhibitor was changed every 24 hours for two days. Timelapse imaging was performed using Nikon AX-R Confocal microscope using Element software. For staining, gels were fixed in 4% formaldehyde for 20 minutes, followed by PBS wash 3 times. Gels were incubated in 1% Triton-X 100 for 10 minutes and washed with PBS for 30 minutes. Next, gels were blocked in 5% BSA solution for 1 hour at room temperature, followed by primary antibody (**Table S1**) incubation overnight at 4°C. Gels were washed 3 times with PBS containing 0.1% TWEEN 20. Gels were then incubated with a conjugated phalloidin probe and secondary antibody (**Table S1**) for 2 hours at room temperature. Gels were washed 3 times with PBS containing 0.1% TWEEN 20 and stored in unsupplemented PBS until imaging (Nikon AX-R Confocal microscope). Timelapse imaging was quantified at a depth of 100 μm, originating from the gel surface using TrackMate (Ershov *et al*, 2022; Tinevez *et al*, 2017) to quantify cell speed along X, Y and Z axes. Images were taken at the center of the gel, total of 6-7 gels from each group were used for analysis with total of 134 cells analyzed. iPericytes in the representative images were pseudo-colored green for better visual distinction from the vascular networks labeled in magenta. The total volume of migrated pericytes in the gel was quantified using Nikon NIS-Elements software. Total of 4 gels from each group were used for analysis with three fields of view for each sample.

### Transwell permeability and transendothelial electrical resistance (TEER) assay

Cells were seeded on type I collagen-coated 24-well Transwell inserts with a 0.4 µM pore size (Corning 3470). After two days, VEGFR2 inhibitor or vehicle control was added to the well for 24 hours. Both assays were conducted 24 hours after the treatment. For the permeability assay, after aspirating the media, cells were incubated with transport buffer containing HBSS (Thermo Fisher Scientific), HEPES (Thermo Fisher Scientific) and 0.1% of platelet poor human serum (Millipore Sigma) for 30 minutes at 37°C. Thereafter, 70 kDa FITC-dextran (250 μg/mL) was applied on the apical side of the Transwell. Samples were collected after 30 minutes to be analyzed with a plate reader at an excitation of 490nm and emission of 520nm. The FITC-dextran concentration was calculated with a standard curve. Relative fluorescence intensity was calculated by normalizing to the average reading of the vehicle control group. For the TEER assay, an EVOM3 system (World Precision Instruments) was used to measure the resistance (Ω). Relative TEER values were calculated by normalizing to average reading of the vehicle control group.

### Retina development model and intravitreal injection

C57BL/6J (Jackson Laboratory) mice were used for all experiments. On P8, female or male mice were intravitreally injected with 0.5 µM of ZM323881 or 4 µM of Tivozanib diluted in PBS in one eye using a microinjector. The contralateral eye was injected with PBS as vehicle control. All studies were conducted under the Duke University Institutional Animal Care and Use Committee-approved animal protocol A070-22-04.

### Oxygen induced retinopathy mouse model and intravitreal injection

We followed a well-established OIR protocol (Cho *et al*., 2020). C57BL/6J (Jackson Laboratory) mice were used for all experiments. Mice were subjected to 75% oxygen from P7 to P12 (BioSpherix ProOx 360). On P12, female or male mice were intravitreally injected with 0.5 µM of ZM323881 or 4 µM of Tivozanib diluted in PBS in one eye by using a microinjector. The contralateral eye was injected with PBS as vehicle control. All studies were conducted under the Duke University Institutional Animal Care and Use Committee-approved animal protocol A070-22-04.

#### Retinal vascular permeability assay in the OIR mouse model

Retinal vascular permeability was measured 5 days after intravitreal injection detailed above. Pups were anesthetized with isoflurane, and 4 kDa FITC-dextran (25 mg/mL) was injected into the retro-orbital venous sinus. Pups were euthanized 30 mins later, eyes were enucleated and immediately fixed in 4% PFA for 10 minutes. The cornea, lens, and retina were removed from the eye cup using dissection forceps and micro scissors under a dissecting scope. Retinas were stained and imaged as detailed below. For quantification, FITC-dextran was extracted from the retina by incubating the tissue in 200 µL N,N-Dimethylformamide (Sigma-Aldrich) overnight at 60°C. Fluorescence intensity was measured with a plate reader.

### Retina flat mount and Immunofluorescence staining

Eyes from P17 pups (OIR or healthy) were enucleated and immediately fixed in 4% PFA for 30 minutes. The cornea, leans, and retina were removed from the eye cup using dissection forceps and micro scissors under a dissecting scope. The isolated retinas were blocked for 1.5 hours using 10% normal goat serum containing 0.3% Triton X-100 in PBS. The retinas were then incubated with DyLight 594-conjugated Griffonia Simplicifolia Lectin I (GSL I) Isolectin B4 (Vector Laboratories) and Alexa Fluor-conjugated Anti-NG2 overnight at 4°C in 5% normal goat serum containing 0.3% Triton X-100 in PBS. Retinas were washed and mounted on the slide using Fluoromount-G (Invitrogen). Images of the samples were obtained using a Nikon AX-R Confocal microscope using Element software. Avascular area and neovascular tufts were quantified by comparing the number of pixels in the area of VO (VO%) or neovascular tufts (NV%) with the total number of pixels in the retina. Mice with body weight lower than 5g on the day of tissue harvest (P17) were excluded from analysis.

Retina cross sections were prepared by fixing the eyes from P17 pups in 4% PFA on ice for 10 minutes and embedding into OCT overnight in the -20°C. Sections were cut along the direction paralleling to the sagittal plane of the eyecup on a cryostat at 8 µm. Immunofluorescence staining was conducted by permeabilizing and blocking the cryosections with 5% normal goat serum containing 0.05% TWEEN 20 in PBS follow by incubation of DyLight 594-conjugated Griffonia Simplicifolia Lectin I (GSL I) Isolectin B4 (Vector Laboratories) overnight at 4°C in the blocking buffer. Slides were washed and incubated with DAPI for 10 minutes the next day. Samples were mounted and imaged on a Nikon AX-R Confocal microscope using Element software.

### Statistics

Biological replicates of in vitro assays are indicated as N and technical replicates are indicated as n. For *in vivo* experiments, mice number in each experiment is indicated as n. Details of replicates for each experiment can be found in the figure legends. All bar graphs represent means±SD. A two-tailed unpaired or paired Student’s t-test were used to determine significance (GraphPad Prism 10). Significance levels were set at *p* > 0.05, *\*p* ≤ 0.05, *\*\*p* ≤ 0.01, *\*\*\*p* ≤ 0.001 and *\*\*\*\*p* ≤ 0.0001.

## Acknowledgments

We thank Dr. Clay Rouse for assisting with animal studies and Dr. Xi Chen for insightful input on the manuscript.

## Author contributions

Conceptualization: YL, BLM, SG

Investigation: YL, EW, BLM, LR

Supervision: SG

Writing-original draft: YL, EW, SG

Writing-review & editing: YL, EW, BLM, LR, SG

## Data and materials availability

All data needed to evaluate the conclusions in the paper are present in the paper and/or the Supplementary Materials. Additional data related to this paper may be requested from the authors.

